# propr: An R-package for Identifying Proportionally Abundant Features Using Compositional Data Analysis

**DOI:** 10.1101/104935

**Authors:** Thomas Quinn, Mark F. Richardson, David Lovell, Tamsyn Crowley

## Abstract

In the life sciences, many assays measure only the relative abundances of components for each sample. These data, called compositional data, require special handling in order to avoid misleading conclusions. For example, in the case of correlation, treating relative data like absolute data can lead to the discovery of falsely positive associations. Recently, researchers have proposed proportionality as a valid alternative to correlation for calculating pairwise association in relative data. Although the question of how to best measure proportionality remains open, we present here a computationally efficient R package that implements two proposed measures of proportionality. In an effort to advance the understanding and application of proportionality analysis, we review the mathematics behind proportionality, demonstrate its application to genomic data, and discuss some ongoing challenges in the analysis of relative abundance data.

## Introduction

Advances in the technology used to assay biological systems have led to a rapid increase in the amount of data generated. However, there has not yet emerged a clear consensus over the best methodological approach for dealing with these data. In fact, some of the most commonly used methods fundamentally ignore the underlying nature of the data studied. Biological count data, such as those produced by high-throughput sequencing, belong within the domain of relative data. Other examples of relative data include data generated by RNA-sequencing (RNA-Seq), chromatin immunoprecipitation sequencing (ChIP-Seq), Methyl-Capture sequencing, 16S amplicon-sequencing, and metabolomics. Methods that wrongly assume absolute data yield erroneous results when applied to relative data, as evidenced by the spurious correlations that arise during the analysis of gene expression data [10].

Genomic count data (i.e., gene expression data from RNA-Seq), as a type of compositional data, have two key geometric properties. First, the total sum of all component values (sometimes called the library size) is an artifact of the sampling procedure [17]. Any number of factors, such as technical variability or differences in experiment-specific abundance, can impact the library size. Second, the distance between component values is only meaningful proportionally (e.g., the difference between 100 and 200 counts carries the same information as the difference between 1000 and 2000 counts) [17]. Simply dividing count data by the total library size does not address the systematic biases in RNA-Seq data. Although several normalization strategies exist, it is concerning that the choice of method can drastically change the number and identity of genes reported as differentially expressed [9]. This sensitivity to normalization holds true for other modes of compositional data as well [16]. Moreover, some methods, such as trimmed mean of M (TMM) normalization, give different results depending on how lowly expressed genes get removed from the data [9].

By not addressing the compositional nature of genomic count data, investigators implicitly assume that absolute differences between counts have meaning [5]. That is, they assume the data exist in real Euclidean space [17]. This may explain why there has emerged a number of hyper-parameterized, assay-specific normalization methods used in the calculation of differential abundance that fail to generalize to data produced by other assays [5]. Meanwhile, methods that accommodate compositional data, such as the ALDEx2 package for R, offer a unified way to compute differential abundance regardless of the data source [5]. However, the ALDEx2 package lacks an interface for calculating proportionality or any other compositionally valid measure of association.

The compositional nature of genomic count data also has considerable importance when measuring correlation because correlation presupposes that the data are measured as absolute abundances. Rather, many of the data studied in genomics are measured as relative abundances, making correlation a statistically invalid choice when working with genomic count data [10]. As Pearson warned in 1896, correlation gives spurious results when applied to relative data: i.e., given three statistically independent variables, X, Y, and Z, the ratios X/Z and Y/Z will correlate with one another because of their shared denominator [12]. If we consider that Z may represent, for example, the library size, we see how two uncorrelated features, X and Y, may appear correlated even when they are not. We emphasize that the topic of spurious correlation is not merely a pedantic footnote. When applied to real biological data, correlation can lead to wrong conclusions [6] [10]. For example, one might incorrectly conclude that there exists a coordinated regulation among a module of transcriptionally independent genes.

As an alternative to correlation, proportionality is a measure of dependence that is valid for compositional data [10] [4]. Borrowing from compositional data analysis (CoDA) principals, this approach uses a log-ratio transformation (*LR) of the original feature vectors in order to transpose the data from a simplex into real Euclidean space [1] [17]. These transformed abundances then provide a substrate for calculating the *log-ratio* variance (VLR), defined as the variance of the ratio of two log-ratio transformed feature vectors (var (*LR (*X*)=*LR (*Y*))) [1]. As a compositionally coherent measure, the VLR does not change whether applied to relative values or to their absolute equivalent. However, the VLR lacks a scale that would otherwise make it possible to compare dependency across multiple feature pairs. In essence, what we call proportionality is a modification to the VLR that establishes scale.

The propr package, now available through the Comprehensive R Archive Network (CRAN) [13], implements two measures of proportionality, *ϕ* [10] and *ρ_p_* [4], defined formally in the next section. As an alternative to correlation, proportionality avoids the majority of the spurious results that come with correlation misuse, having a certain robustness in the setting of compositional data [10]. Between *ϕ* and *ρ_p_*, *ρ_p_* has one benefit in that it provides a naturally symmetric result. However, we can make *ϕ* naturally symmetric too by slightly altering its definition, as shown in the next section. Otherwise, *ρ*_p_ exists on a scale from [−1; 1], reinforcing its analogy to correlation, while *ϕ* exists on a scale from [0; ∞), reinforcing its analogy to dissimilarity. Yet, it is the conceptual and mathematical link between *ϕ* and *ρ_p_* metrics that allows us to present the propr package as a single portal to proportionality analysis.

## Design and Implementation

This software tool provides a programmatic framework for calculating two measures of proportionality in the R language. Proportionality, as analogous to, but distinct from, correlation, measures the dependence between two feature vectors. Unlike correlation, which can yield spurious results in the setting of relative data, proportionality borrows from the branch of mathematics known as compositional data analysis (CoDA) to offer a robust alternative for relative data types. This package includes two principal functions, phit and perb, that calculate the proportionality metrics *ϕ* and *ρ_p_*, respectively [10] [4].

Consider a matrix of D values measured across N samples subjected to one of any number of binary or continuous events M. The events M might represent case-control status, treatment status, treatment dose, or time. Given a dataset with N rows and D columns, the functions phit and perb return a matrix of *D^2^* elements relating each combination of two vectors, *A_i_* and *A_j_*, according to the following definitions:

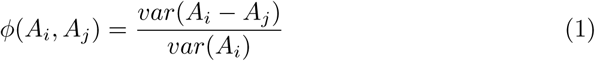

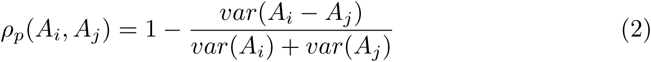

In addition, we include the phis function to calcualate *ϕ_s_*, a naturally symmetric variant of *ϕ*. Interestingly, *ϕ_s_*, relates to *ρ_p_* by a monotonic function [4].

This measure of proportionality relates the variance of the log-ratio (VLR) to the variance of the log-product (VLP), according to the following definition:

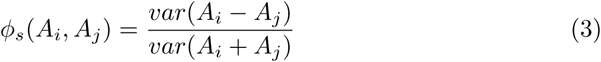

We refer the reader to the supplemental appendix for a demonstration of how these measures of proportionality relate to the slope, β, and the Pearson correlation coefficient, r, of pairwise log-transformed data. This appendix also illustrates how *ϕ*, *ϕ_s_*, and *ρ_p_* relate to one another S1 Appendix.

Above, we use *A_i_* and *A_j_* to denote a log-ratio transformation of the sample vectors *X_i_* and *X_j_*. By default, this package uses an implementation for the centered log-ratio transformation (clr), defined as:

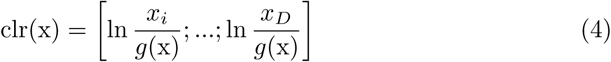

In addition, we include an implementation for the additive log-ratio transformation (alr), defined as:

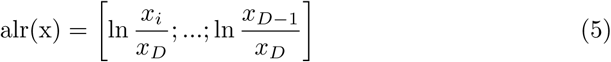

In contrast to clr, alr uses another feature to transform the original sample vectors. When used in conjunction with an *a priori* known unchanged reference, alr effectively back-calculates the absolute counts from the relative components. By specifying, for example, a house-keeping gene or an experimentally fixed variable, the investigator can possibly achieve a more accurate measure of dependence than through clr [4]. The user can toggle alr transformation in lieu of the default clr transformation by supplying the name of the unchanged reference to the ivar argument of the perb or phis function. In either case, we wish to alert the reader that log-ratio transformation, by its nature, require non-zero elements in the data matrix. As such, any log-ratio analysis must first address any zeros. Yet, how best to do this remains an open question and a topic of active research [11]. For simplicity, propr automatically replaces all zero values with 1 prior to log-ratio transformation, corresponding to the multiplicative replacement strategy [11].

The R language, despite its widespread popularity, suffers from poor performance when scaling to big data. Its “copy-on-modify” behavior, whereby each modification of an object creates a duplication of that object, along with slow for-loops, makes it an imperfect choice for computationally expensive tasks such as the one here. Therefore, in order to speed up the run time and reduce RAM overhead, we harness the Rcpp package to draft the computationally expensive portions of this tool in C++ [3].

This package also offers a number of tools to visualize proportionality when working with high-dimensional data. We provide extensive documentation for these plotting methods in the package vignette, “Understanding RNA-Seq Data through Proportionality Analysis”, hosted with the package on CRAN [13]. Among these tools are those used to generate the figures included in the next section.

## Results

As a use case, we re-analyze the raw RNA-Seq counts from an already published study on cane toad (*Rhinella marina*) evolution and adaptation [15]. Sugar cane farmers introduced cane toads to Australia in 1935 as a cane beetle pest control measure, but these toads quickly became invasive. This event now serves as a notable example of failed biological control. Initially introduced into northeastern Australia (Queensland, QLD), cane toads have since spread westwards across the continent to Western Australia (WA) [15]. This dataset contains muscle tissue RNA transcript counts for 20 toads sampled from two regions (10 per region) in the wild. The two regions sampled, which we will treat as the experimental groups, include the long colonized site of introduction in QLD and the front of the range expansion in WA [15]. In this analysis, we want to understand the differences in gene expression between the established and expanding populations. By demonstrating propr on public data, we provide a reproducible example of how proportionality analysis can converge on an established biological narrative. The reader can find these data bundled with the release of the package on CRAN [13].

We begin by constructing the proportionality matrix using all 57,580 transcript counts, yielding an *N*^2^ matrix 24.7 Gb in size. To minimize the number of lowly expressed transcripts included in the final result, we subset the matrix to include only those transcripts with at least 10 counts in at least 10 samples. By removing the features at this stage, we can exploit a computational trick to calculate proportionality and filter simultaneously, reducing the required RAM to only 5 Gb without altering the resultant matrix. Next, in the absence of a hypothesis testing framework, we arbitrarily select those “highly proportional” transcripts with *ρ_p_ > 0:99*. When plotting the pairwise log-ratio transformed abundance for these “highly proportional” transcript pairs, a smear of straight diagonal lines confirms that the feature pairs indexed as proportional actually show proportional abundance (Fig 1).

The procedure used to parse through the proportionality matrix now depends on the experimental question. Here, we wish to identify a highly proportional transcript module that happens to show differential abundance across the experimental groups. In this example, we take an unsupervised approach by hierarchically clustering the highly proportional feature pairs based on the matrix 1 abs(*ρ_p_ > 0:99*). We note here that one could instead cluster using *ϕ_s_* directly. When clustering, we call two features co-clustered if they belong to the cluster after cutting the dendrogram. Then, we project the pairs across two axes of variance, the variance of the *log-ratio* (VLR) and *variance of the log sums* (VLS) such that 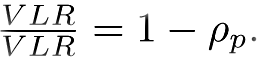 In this formula, we see that as VLR approaches 0, *ρ_p_* approaches 1. Meanwhile, the VLS, the sum of the individual variances of two features in that pair, adjusts the rate of this limit. Since we would expect a differentially expressed module to have a low VLR and a high VLS, we prioritize pairs in co-cluster 3 for subsequent analysis (Fig 2).

**Fig 1.**
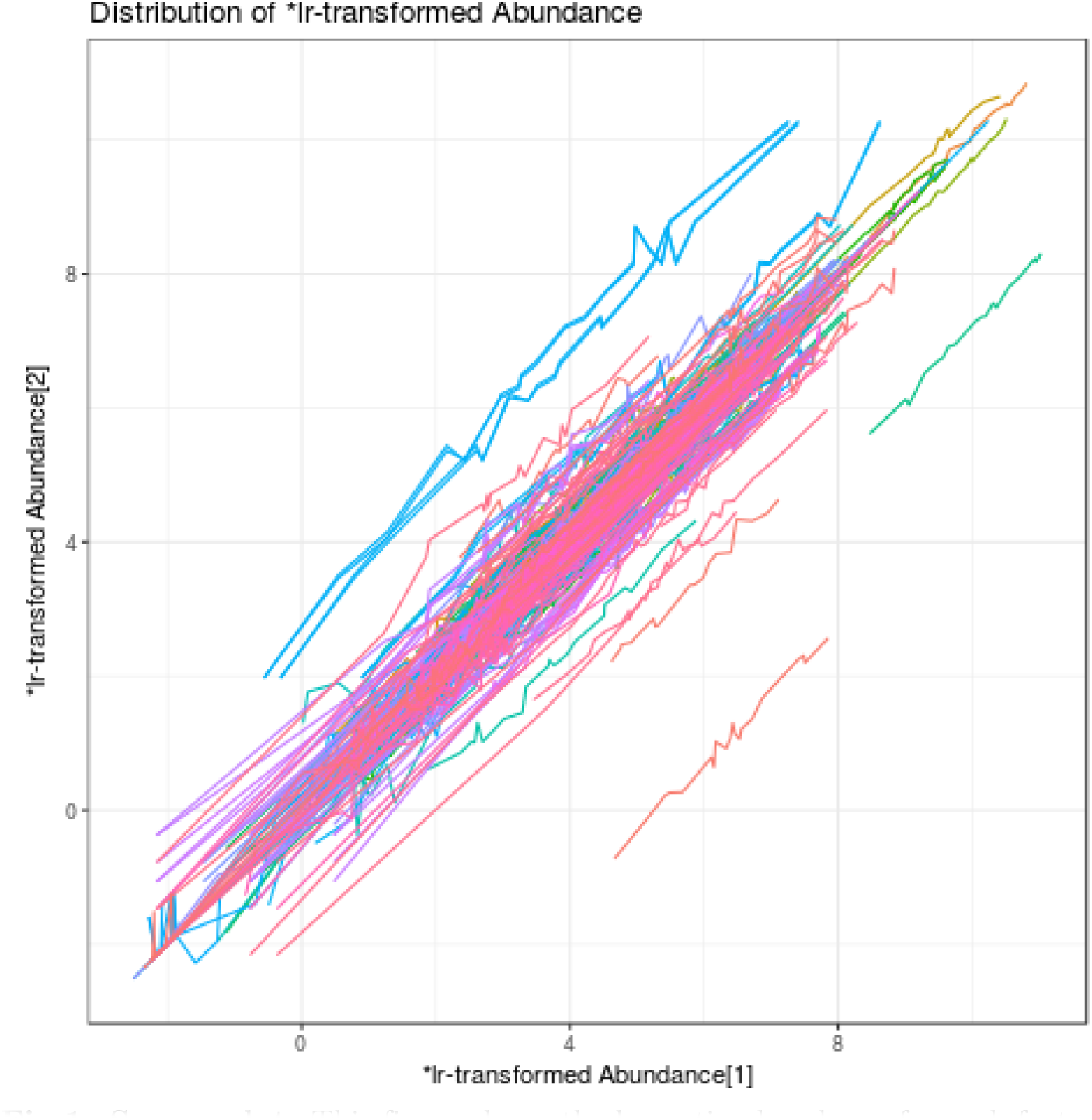
Smear plot. This figure shows the log-ratio abundance for each feature belonging to pairs index as highly proportional (*ρ_p_ > 0:99*). A smear of straight diagonal lines confirms that the feature pairs indexed as proportional actually show proportional abu e. In other words, large deviations from *y = x* indicate small values of 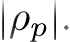 Figure produced using the smear function in propr.

Co-clusters containing feature pairs with a low VLR and a high VLS have the potential to explain differences between the experimental groups. However, the high VLS may not necessarily have anything to do with the experimental condition. For example, this co-cluster might instead include highly proportional features that show wide individual feature variance due to batch effects. For the cane toad data, however, the experimental condition does indeed seem to drive the high individual feature variances in the module, as evidenced by the near perfect group separation when visualizing the first two components of a principal components analysis (PCA) (Fig 3). Note that this plot calculates PCA using the log-ratio transformed data, making it a statistically valid choice for compositional data [7]. The separation between groups achieved here compares to that reported in the original publication, which used features selected by the edgeR package [15]. In addition, gene set enrichment analysis of the gene ontologies for co-cluster 3 (S1 Table) shows an enrichment for similar molecular functions as those highlighted in the original publication [15]. Although the exact gene ontology terms differ by method, our approach yields a constellation of macromolecular metabolic terms that suggests a common story: “metabolic enzymes are overwhelmingly upregulated at the invasion front” [15], interpreted to mean that WA cane toads may experience more environmental stress than those from QLD [15]. Yet, while agreeing here, proportionality analysis offers an additional benefit in that it provides a layer of information on pairwise associations, all without requiring any kind of normalization.

**Fig 2.**
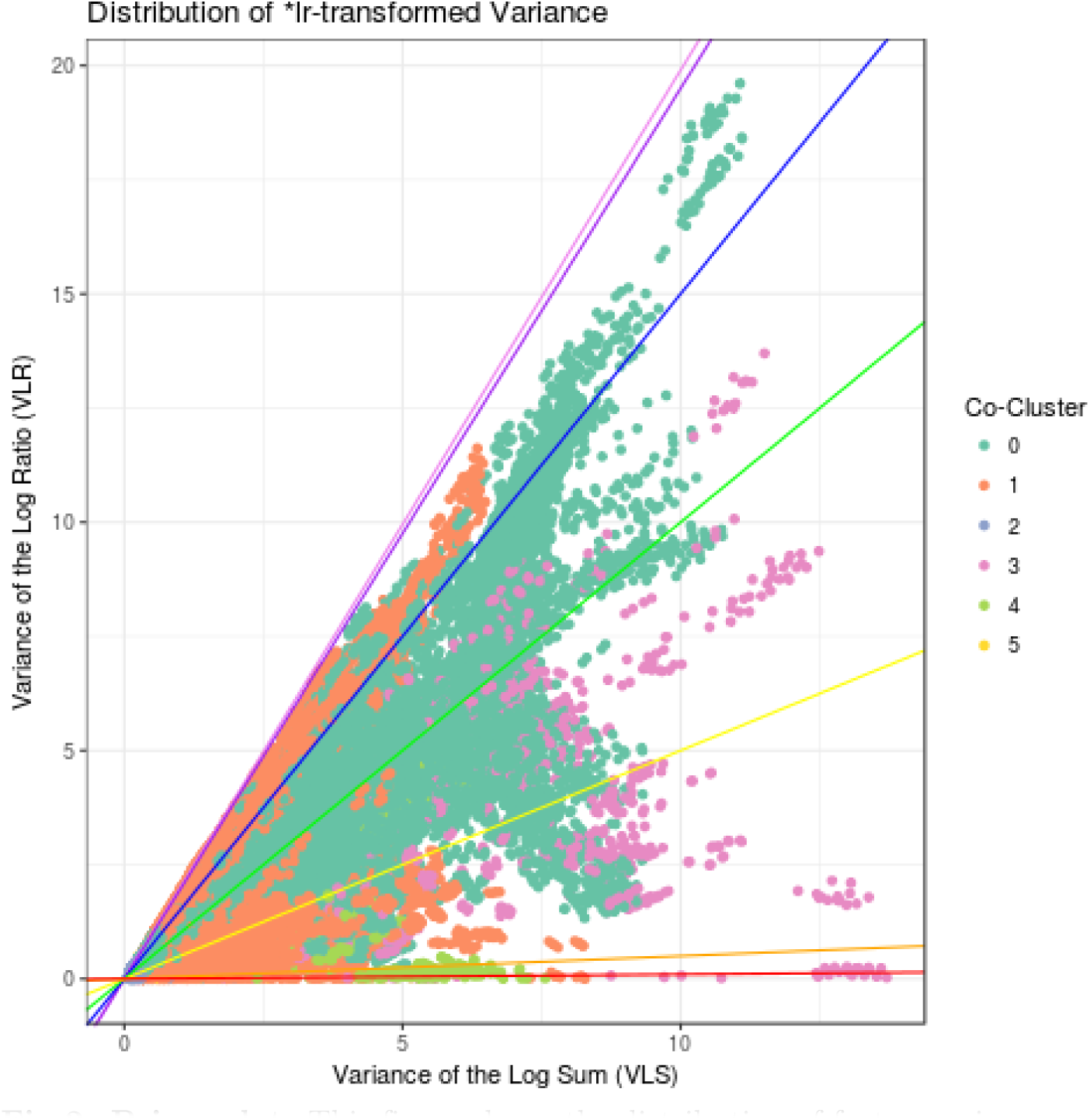
Prism plot. This figure shows the distribution of feature pairs according to the variance of the log-ratio (VLR) and the variance of the log sums (VLS) for all pairs in which at least one of the features participates in at least one highly proportional (*ρ_p_ >* 0:99) pair. If both features in a pair belong to the same cluster, they receive a non-zero color code. Clusters created hierarchically based on the matrix 1 abs (*ρ_p_*). Figure produced using the prism function in propr.

**Fig 3.**
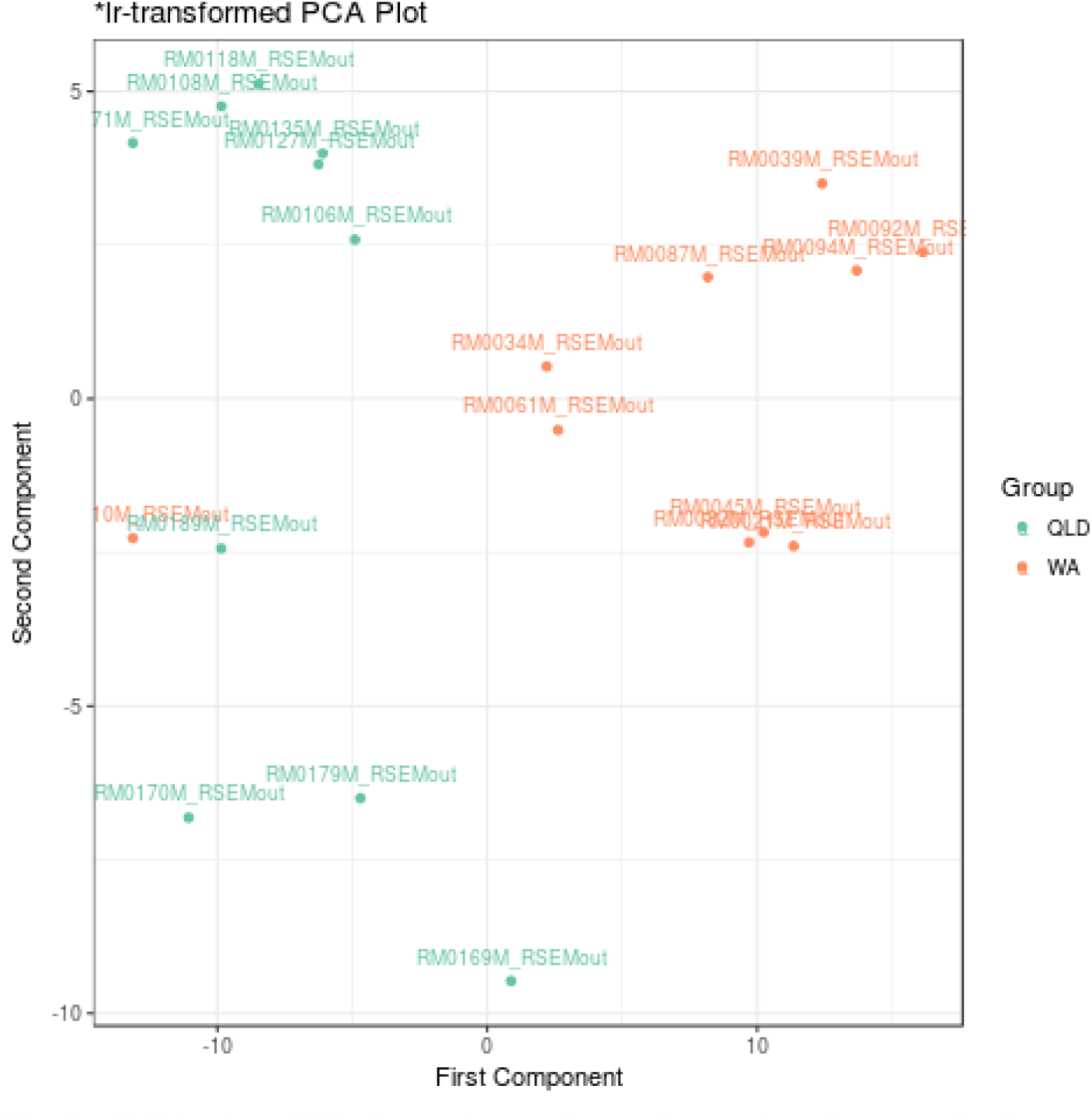
PCA plot. This figure shows all samples projected across the first two components of a principal components analysis (PCA), calculated using the log-ratio transformed data. This plot colors samples based on the experimental group. Figure produced using the pca function in propr.

We refer the reader to the supplementary materials for a script that contains everything required to reproduce this analysis (S1 File).

## Availability and Future Directions

The propr package for R, now available on CRAN [13], provides a fast implementation of two measures of proportionality previously shown to rectify the issue of spurious correlation in the setting of compositional data [10] [4]. These implementations offer a valid alternative to correlation that can accurately identify feature dependence from relative data [4]. By using the Rcpp package to draft the computationally expensive code in C++, the propr package achieves greater performance than possible in the native R environment without altering the front-end user experience [3]. In this way, proportionality analysis executes nearly as fast as base R correlation while retaining a simple programming interface. Yet, proportionality analysis is not without limitations.

First, unlike the *log-ratio variance* (VLR), the values of Φ and *ρ_p_* will change in the setting of missing feature data. Since biological assays rarely, if ever, capture all possible feature information, this property makes the centered log-ratio transformation (clr) a sub-optimal choice. The additive log-ratio transformation (alr), which allows the user to scale their data by a feature with a priori known fixed abundance, such as a house-keeping gene or an experimentally fixed variable (e.g., a ThermoFisher ERCC synthetic RNA “spike-in” [8]), may provide a superior alternative. In contrast to clr, proportionality calculated with alr does not change with missing feature data because it effectively back-calculates the absolute feature abundance. Still, clr remains sufficient when most features remain unchanged across samples, an assumption also built into the trimmed mean of M (TMM) normalization used with RNA-Seq data [14] [4]. Moreover, errors associated with clr transformation consist almost entirely of false negatives, rather than false positives, making it suitable for many scientific applications [4].

Second, proportionality analysis, by nature of the log-ratio transformation, fails in the presence of zero values. As such, proportionality analysis inherits the zero replacement controversy prominent in the compositional data analysis literature. By default, propr replaces all zero values with 1. If analysts wish to explore other approaches to zero replacement{including conducting analyses of the sensitivity of results to zero imputation{they can so by manipulating the data prior to running routines in propr. We stress that, depending on the number of zero values in the data, replacement of zeros can have a major impact on the conclusions drawn from any log-ratio compositional data analysis. Analysts should carefully examine their results to understand the extent to which they depend on the zero replacement strategy chosen.

Third, biological count data do not exist as true compositional data, but rather as a kind of “count-compositional” data, whereby small non-zero counts pose a unique challenge to analysis. This follows from how the log-ratio methods of compositional data analysis assume data to consist of *D* positive, real-valued components (i.e., a sample space of 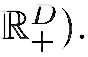. Instead, count-compositional data consist of D non-negative, *integer-valued* components (i.e., a sample space of 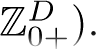 The smaller the counts, the more noticeable the discretization of the count-compositional data becomes. Likewise, the impact of sampling variation becomes more noticeable as well: additive variation affects the relative abundance of small counts more than large counts. Put succinctly, the difference between 1 count and 2 counts does not carry the same information as the difference between 1000 counts and 2000 counts. In practice, this means that a minor variation in small counts can have a major impact on the conclusions drawn from any log-ratio compositional data analysis. Like with zero replacement, analysts should carefully examine their results to understand this sensitivity. We note that prior work has found substantial differences in how test statistics handle low-count genes in RNA-Seq [2]. For the purposes of demonstrating propr, we avoid this issue by altogether removing from analysis any component with a predominance of low counts.

Finally, proportionality analysis currently lacks a hypothesis testing framework. Its distribution of heteroscedastic variance makes it unsuited for the z-transformation used to calculate the variance of the correlation statistic. More work is needed to create a framework for applying statistical tests to proportionality analysis. However, despite these limitations, we believe the propr package comes a long way in improving the accessibility of proportionality analysis to researchers. We hope many biological investigations will benefit from this alternative to correlation.

## Supporting Information

**S1 Appendix. Examination of proportionality**. This document lays out how the measures of proportionality relate to the slope, *β*, and the Pearson correlation coefficient, r, of pairwise log-transformed data, as well as how these measures relate to one another.

**S1 File. Cane toad analysis script**. This R script reproduces the results and figures presented in this paper.

**S2 File. Gene ontology (GO) data object**. This R object provides the gene ontology terms used by S1 File to produce S1 Table. Function annotations sourced from the Trinotate pipeline (http://trinotate.sourceforget.net), as described in the supplementary methods from [15].

**S1 Table. Gene ontology (GO) table**. This table contains the gene ontology terms selected by gene set enrichment analysis of the highly proportional (*ρ_p_ >* 0:99) gene pairs.

